# Preponderant Myosin Super-Relaxed State In Skeletal Muscle From Endurance Athletes

**DOI:** 10.1101/2022.09.23.509202

**Authors:** Christopher T. A. Lewis, Lee Tabrizian, Joachim Nielsen, Jenni Laitila, Thomas N. Beck, Per Aagaard, Rune Hokken, Simon Laugesen, Arthur Ingersen, Jesper L. Andersen, Casper Soendenbroe, Jørn W. Helge, Flemming Dela, Steen Larsen, Ronni E. Sahl, Tue Rømer, Mikkel T. Hansen, Jacob Frandsen, Charlotte Suetta, Julien Ochala

## Abstract

It has recently been established that myosin, the molecular motor protein, is able to exist in two conformations in relaxed skeletal muscle. These conformations are known as super-relaxed (SRX) and disordered-relaxed (DRX) states and are finely balanced to optimize skeletal muscle metabolism. Indeed, SRX myosins are thought to have a 10-fold reduction in ATP turnover compared to DRX myosins. Here, we investigated whether chronic physical activity in humans would be associated with changes in the proportions of SRX and DRX skeletal myosins. For that, we isolated muscle fibres from various athletic and sedentary populations and ran a loaded Mant-ATP chase protocol. We observed that, in endurance-trained athletes, the amounts of myosin molecules in the SRX state was significantly greater than in age-matched sedentary individuals or than in strength athletes. To further assess whether this change would have an impact on the potency of a SRX-inducing pharmacological compound, Mavacamten, we performed similar analyses as above with and without the drug in muscle fibres from endurance athletes. Surprisingly, we found that 0.3 μM of Mavacamten had only marginal effects. Altogether, our results indicate that chronic endurance training-status influences resting skeletal myosin conformations, and Mavacamten potency. Our findings also emphasize that environmental stimuli such as exercise can re-wire the molecular metabolism of human skeletal muscle through myosin.

**Summary:** Lewis *et al*., investigate how training-status influences myosin conformations involved in the resting metabolism of skeletal muscle. They find that, in endurance-trained athletes, skeletal myosin preferentially adopts an energy-saving conformation known as super-relaxed state, lowering the metabolic rate and affecting the potency of a super-relaxed state-inducing drug, Mavacamten.

## Introduction

The role of myosin II as a vital motor molecule in the contraction of striated muscle has been the subject of intense research over recent decades (Trivedi et al., 2018). What remains unclear is how myosin can regulate muscle function at rest. Myosin was once thought to be a passive molecule in relaxed muscle however, this paradigm changed in 2010 upon the discovery that myosin can exist in two different resting conformations (Stewart et al., 2010). These different conformations, or states, are the disordered-relaxed (DRX) and super-relaxed (SRX) states (Cooke, 2011). These states differ by the relative spatial positioning of myosin heads between thick and thin filaments (Toepfer et al., 2020). In the DRX state, one of the two myosin heads is free to sway within the inter-filament space of the sarcomere and thus both the actin-binding and ATP-binding motifs on this myosin head are sterically open (Alamo et al., 2016, Alamo et al., 2017, Lee et al., 2018). In the SRX state, both myosin heads are folded back onto the thick-filament backbone and, therefore, the ATP-binding motif on both myosin heads in this molecule are sterically inhibited (Naber et al., 2011, Alamo et al., 2016, McNamara et al., 2015). This inhibition of the ATP-binding motif on the myosin head causes the substantial functional difference between the DRX and SRX conformations; with the SRX conformation demonstrating a ten-fold reduction in ATP turnover compared to the DRX conformation (Cooke, 2011, Toepfer et al., 2020, Schmid and Toepfer, 2021). This key difference between the two resting myosin conformations has led to the theory that SRX may have evolved as an energy saving mechanism to reduce ATP consumption in cardiac and skeletal muscle (Lee et al., 2018, Hooijman et al., 2011).

As the discovery of the SRX state is still relatively novel, little is known about the role of resting myosin conformations in the context of disease or in the adaptation to environmental stimuli. Much of our knowledge of the regulation and functional significance of resting myosin head conformations comes from the discovery that dysregulation of myosin head conformation in cardiac muscle is pathogenic in hypertrophic cardiomyopathy (HCM) (Alamo et al., 2017). HCM is a genetic cardiomyopathy caused by mutations in sarcomeric proteins, including myosin (Debold et al., 2007, Dijk et al., 2009). As a disease it results in myocardial hyper-contractility, hyper-metabolic rate and reduced relaxation, which leads to progressive heart failure and may cause sudden cardiac arrest (Maron and Maron, 2013). It has been observed that in the various mutations that cause HCM, there are significantly higher numbers of DRX myosins in the cardiac muscle of these patients (Schmid and Toepfer, 2021, Toepfer et al., 2020, Toepfer et al., 2019). The discovery of increased DRX myosins in HCM patients was significant as it presented a novel target for the treatment of HCM. This proved to be successful and in 2021 the Federal Drug Agency (FDA) approved, Mavacamten, a first-in-class drug for the treatment of HCM (Olivotto et al., 2020). Mavacamten is an allosteric modulator that inhibits the ATP turnover of myosin heads by stabilizing the SRX state, functionally reducing the number of DRX myosins (Anderson et al., 2018, Gollapudi et al., 2021). As a result, cardiac function is restored in HCM patients (Olivotto et al., 2020). This positive finding has led to the promise that myosin head conformations may be able to be targeted for the successful treatment of other diseases, such as skeletal myopathies. However, no convincing data exists as of yet showing that compounds targeting myosin heads can act upon human skeletal muscle fibres.

To date, the involvement of myosin heads in disease and their dysregulation has all been attributed to genetic mutations in either myosin or neighbouring partners. Therefore, an important question that remains to be addressed is how dynamic these different conformations are in their nature and if they can be influenced by lifestyle changes such as diet or physical activity/training. Given that exercise has vast influence upon the function of skeletal muscle tissue and its molecular determinants, particularly in the context of the metabolic rate, in the present study, we aimed to explore whether training-status in humans would be linked to remodeled proportions of SRX and DRX skeletal myosins. For that, we specifically analysed muscle fibres extracted from muscle biopsy specimens from four distinct groups, sedentary controls, physically active controls, endurance-trained athletes and strength-trained athletes. We initially hypothesized that chronic endurance and strength trainings would both favour SRX myosins in order to conserve energy and thereby optimize energy re-charge rates. Following this, we also hypothesized that Mavacamten, the pharmacological compound presented above promoting the SRX state would be able to further increase the SRX population in myofibres from endurance-trained athletes.

## Materials and Methods

### Preparation of single muscle fibres from skeletal muscle

Human vastus lateralis biopsy specimens were obtained upon informed consent from four different population groups. Two control groups were used in this study. The first group comprised sedentary controls (SC), were untrained young males that completed less than three sessions of cardiopulmonary exercise per week. The second control group were physically active controls (PAC), these were young male subjects who were engaged in leisure exercise, typically completing three or more cardiopulmonary exercise sessions (45-60 min/session) per week (Sahl et al., 2018). Using these two distinct control groups allowed for the observation of potential differences in the myosin head conformation between two groups with only modest differences in their level of physical activity versus extremely active athletes (120-180 min/day). Two contrasting athlete groups were recruited, comprising endurance athletes (EA) and strength athletes (SA). EA subjects were recruited from local groups of very highly-trained endurance cyclists (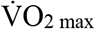 63.0 ± 3.4 mL/kg/min training load 9.1 ± 3.7 h/week) (table 1) (Frandsen et al., 2022). SA subjects were recruited from local weight training clubs. SA subjects were required to have resistance trained for a minimum of three years in the years preceding the study and have the ability to squat at least 1.5-times their own bodyweight (Hokken et al., 2021). These two athlete groups represent individuals that, whilst both highly-trained, have vastly different training regimes and goals. This strategy was chosen to identify if specific training regimes would be able to specifically alter myosin head conformation and affect the metabolism of resting skeletal muscle. The Greater Copenhagen Region science ethical committee approved the use of sedentary controls, physically active controls and endurance athlete biopsies (project IDs: H-16049145, H-15002266). The ethics committee in the Region of Southern Denmark approved the use of strength athlete biopsies (project ID: S-20160116).

**Table 1:** Subject characteristics. Mean subject characteristics for the participants in the studies of which biopsies samples were obtained from. Shown parameters are age (years), height (cm), weight (kg), BMI (kg/m^2^) for all subjects. 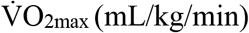 is shown for physically active control and endurance athlete subjects. Data is displayed as mean ± SD. ** = p<0.01 vs SC. Ψ = p<0.0001 vs PAC.

|  | Sedentary Controls (SC) (n=6) | Physically Active Controls (PAC) (n=6) | Endurance Athlete (EA) (n=6) | Strength Athletes (SA) (n=6) |
| --- | --- | --- | --- | --- |
| **Age (Years)** | 31.2 ± 9.4 | 24.3 ± 3.7 | 31.0 ± 4.4 | 24.4 ± 2.6 |
| **Height (cm)** | 178.5 ± 6.0 | 189.3 ± 7.0** | 181.3 ± 4.6 | 180.8 ± 3.8 |
| **Weight (kg)** | 74.2 ± 8.9 | 90.8 ± 9.9** | 77.5 ± 4.9 | 95.4 ± 10.0** |
| **BMI (kg/m<sup>2</sup>)** | 23.3 ± 2.7 | 25.3 ± 2.9 | 23.5 ± 0.5 | 29.3 ± 3.2** |
| **$\dot{V}O_{2max}$ (mL/kg/min)** | N/A | 44.8 ± 4.0 | 63.0 ± 3.4<sup>ψ</sup> | N/A |

### Physical performance testing

For EA 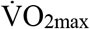 was assessed with breath-by-breath measurements of pulmonary 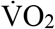 and 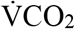 by an online system (COSMED Quark CPET, COSMED, Italy) during a graded exercise test performed on a cycle ergometer. The protocol was adapted from the Achten_35/3_ protocol and described in detail elsewhere (Achten et al., 2002, Frandsen et al., 2022). In brief, the 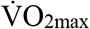 protocol was performed in continuation of a submaximal graded exercise test for determination of substrate oxidation, and 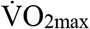 was elicited through workload increments of 35 W every minute until volitional exhaustion.

### Mant-ATP chase experiments

All biopsies were immediately snap frozen in liquid nitrogen and stored at −80°C. At the time of analysis, single muscle fibres were chemically skinned as previously described (Ochala et al., 2021). They were stored at −20°C in a buffer solution containing glycerol and relaxing solution mixed 50/50. As previously published, single myofibres were individually attached to a TEM-grid (Sigma-Aldrich, Grids for transmission electron microscopy, G1403-1VL, UK), which was positioned on a microscopy slide (Thermo Scientific, Superfrost ^®^ Plus, USA) (Ochala et al., 2021). Single nucleotide turnover was measured in flow chambers using fluorescent mant-ATP (250μM) as described previously (Stewart et al., 2010, Ochala et al., 2021). Single muscle fibres with a relaxed sarcomere length of 2.2 μm were incubated in a rigor buffer containing 100 μL mant-ATP buffer for five minutes. Rigor buffer contained 120 mM Potassium acetate, 5 mM Mg acetate, 2.5 mM K2HPO4 (dibasic), 2.5 mM KH2PO4 (monobasic), 50 mM MOPS, 5 mM EGTA, 100mM TCEP and KOH to pH 6.8. Mant-ATP or ATP were added to the rigor buffer to make the Mant-ATP buffer of ATP chase buffer respectively. Following incubation, fibres were positioned in the microscope and chased with 100 μL of rigor buffer containing ATP (4 mM). Fluorescence decays due to the mant-nucleotide dissociation from myosin was monitored over 300s. The excitation wavelength for mant-ATP was 385 nm. All experiments were performed at room temperature, 25°C. Fluorescence images were acquired on a Zeiss AXIO Lab A1 microscope (Carl Zeiss AG, GE). The average fluorescence intensities within a selected region of the fibre were determined using the ImageJ software. Background intensity measurements were also taken. Fibre fluorescence values were determined by subtracting the background intensity from the fibre intensity.

### Preparation of Mavacamten solution

Myosin allosteric modulator Mavacamten was purchased from MyoKardia (Bristol Myers Squibb, US). A stock solution (1 mM) of Mavacamten was first prepared using dimethyl sulfoxide (DMSO). The stock solution was used to achieve a final concentration of 0.3 μM Mavacamten in the buffers for experimental use. The 0.3 μM concentration of Mavacamten was chosen as it represents the IC_50_ value for inhibiting ATPase activity in myosin in assays using bovine and murine myosin (Green et al., 2016).

### Mant-ATP chase experiments using Mavacamten

Mavacamten was added to rigor buffer, Mant-ATP buffer and ATP chase buffer solutions as listed above at a concentration of 0.3 μM. From here, the experiment and solutions were performed the same way using solutions including Mavacamten.

### Mant-ATP chase experiment data analysis

Data was plotted using GraphPad Prism 8, after fitting with a two-phase exponential decay function (figure 1) using the following equation:

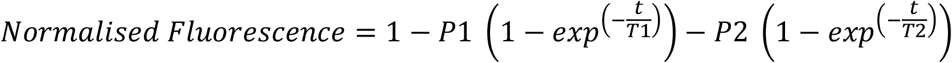

**Figure 1.**
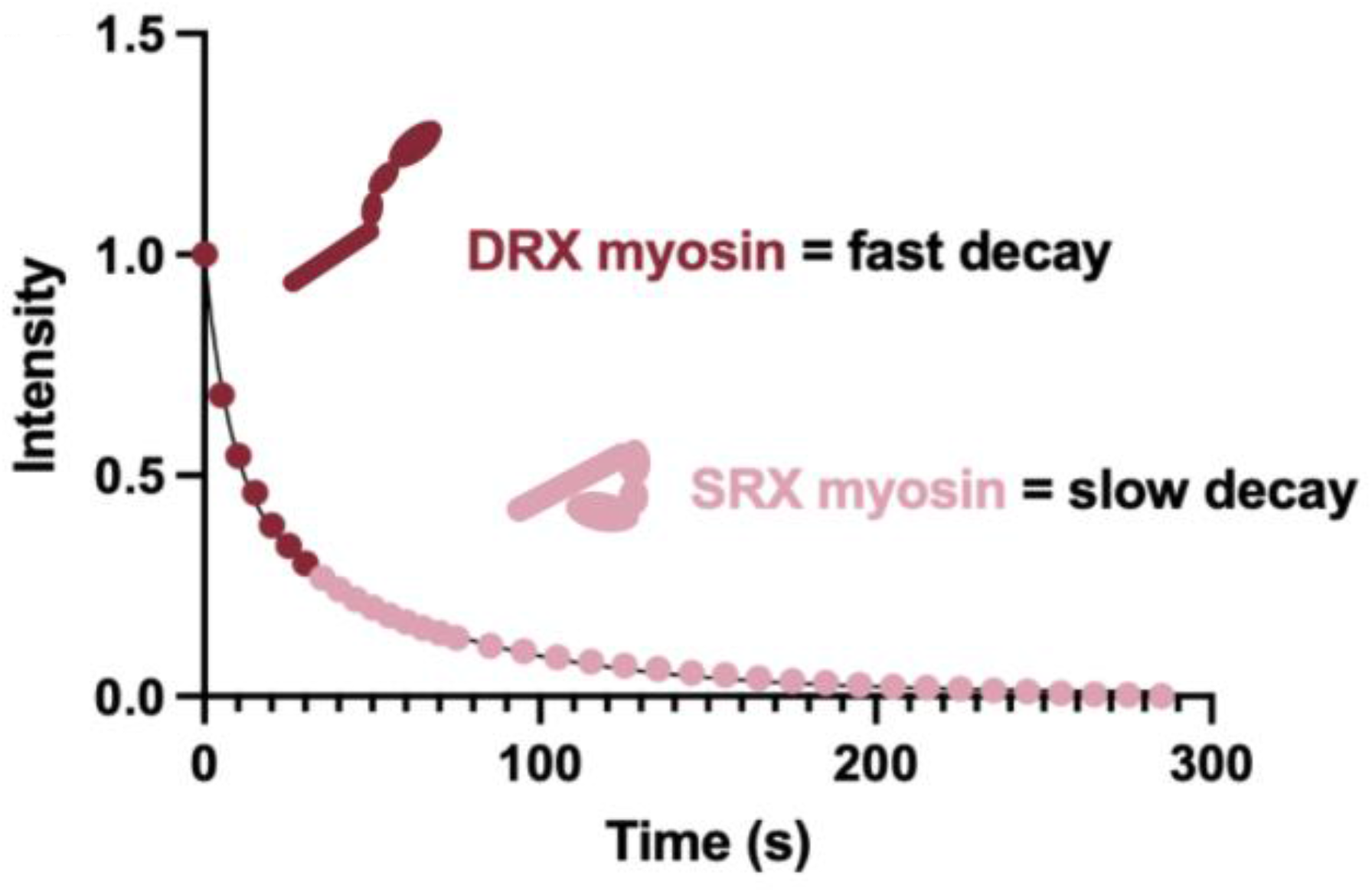
Representative two-phase decay of Mant-ATP chase experiment. This trace demonstrates the decay seen when during the ATP chase period of the Mant-ATP chase experiment. The decrease in fluorescence intensity has two phases, a fast phase (DRX state) and a slow phase (SRX state), measured over a total of 300s.

Where P1 is the amplitude of the rapid decay approximating the disordered-relaxed state (DRX). T1 (seconds) is the time constant for the life of this P1 phase. P2 is the second phase approximating the super-relaxed state (SRX) with the corresponding T2 as the time constant for the life of this P2 phase. As T2 and T2 denote the time constants for their corresponding phase, they are the calculated ATP turnover lifetimes for the corresponding myosin head conformation of which they represent.

### Fibre-typing

After the completion of the Mant-ATP chase experiments using Mavacamten, individual fibres were stained with an anti-myosin slow/type I antibody (IgM isoform: 1:50, DSHB, A4.951). Fibres were then washed in PBS and incubated with a secondary antibody conjugated to Alexa 647 in a goat serum (ThermoScientific, USA, dilution 1:1000). After washing, muscle fibres were mounted in Fluoromount and images taken with a Zeiss AXIO Lab A1 microscope (Carl Zeiss AG, GE, objectives x20 and x10). Positive staining with the myosin β-slow/type I antibody indicated a type I muscle fibre, negative staining indicated a fast/type II muscle fibre.

### Statistical analysis

All measurements for Mant-ATP assays were performed upon 5-6 isolated muscle fibres per individual, fibre data were averaged per individual. Statistical significance was calculated using one-way ANOVA and paired or paired student’s t tests where applicable. p<0.05 was assumed to be significant. Parametric tests were used due to the normal distribution of the data. Data are presented as mean ± SEM in figures and as mean ± SD in table 1 containing subject characteristics. Sample number *n* refers to the number of individuals in each subject group. In paired experiments, data is displayed as the same single muscle fibre before and after Mavacamten treatment, statistical testing was performed upon the mean value of all fibres from each subject. Only slow/type I fibres were analyzed and plotted. In T2 measurements, values of greater than 500s were excluded as outliers. Statistical testing was performed using GraphPad Prism Version 9.3.1 (471) (Insight Partners, USA).

## Results

### Subject information of skeletal muscle biopsies used in study

This study utilized four distinct groups, each with vastly different physical training profiles, several of which have been used in studies published previously (Hokken et al., 2021, Frandsen et al., 2022, Sahl et al., 2018). Age, height, weight, and body mass index (BMI) for all four groups as well as 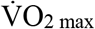 max values for physically active controls and endurance athletes are presented in Table 1. Importantly for this study, none of the subjects were overweight despite a considerable heterogeneity in level of physical activity. Hence, BMI for the SC, PAC and EA groups were within a normal range (23.3 ± 2.7, 25.3 ± 2.9 and 23.5 ± 0.5 kg.m^-2^ respectively) whereas SA had a significantly higher BMI (29.3 ± 3.2, p<0.05) due to a higher lean mass (Westcott, 2012). Otherwise, the 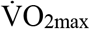 of the EA group was significantly higher than the PAC group (63.0 ± 3.4 vs 44.8 ± 4.0 mL/min/kg, p<0.05), underlining the highly-trained level of the EA group.

### Higher level of SRX myosin heads in the skeletal muscle fibres of endurance athletes

To test our initial hypothesis of a higher proportion of myosin heads in the SRX state in both EA and SA groups, we ran a loaded Mant-ATP chase protocol using extracted myofibres from all four subject groups (P1 and P2 values). Interestingly, the amount of DRX myosins was only significantly lower in the EA group when compared to SC, PAC and SA groups (Figure 2a, b). These changes were matched by equally significant higher levels of SRX myosins for EA when compared to SC, PAC and SA groups (Figure 2c). This result is partially in line with our hypothesis eluding the fact that the regulation of skeletal myosin resting conformations may be influenced in a training-specific manner. Using the same data obtained from our loaded Mant-ATP experiments, we next determined the ATP turnover times of both DRX and SRX myosins of all four groups (T1 and T2 values). We did not observe any significant difference in the ATP turnover time of DRX myosins (Figure 2d). Nevertheless, we found that the ATP turnover time of SRX myosins was significantly greater in the SA group when compared to EA group only (not the other groups) (Figure 2e). Hence, the physiological relevance of such particular finding remains to be determined.

**Figure 2.**
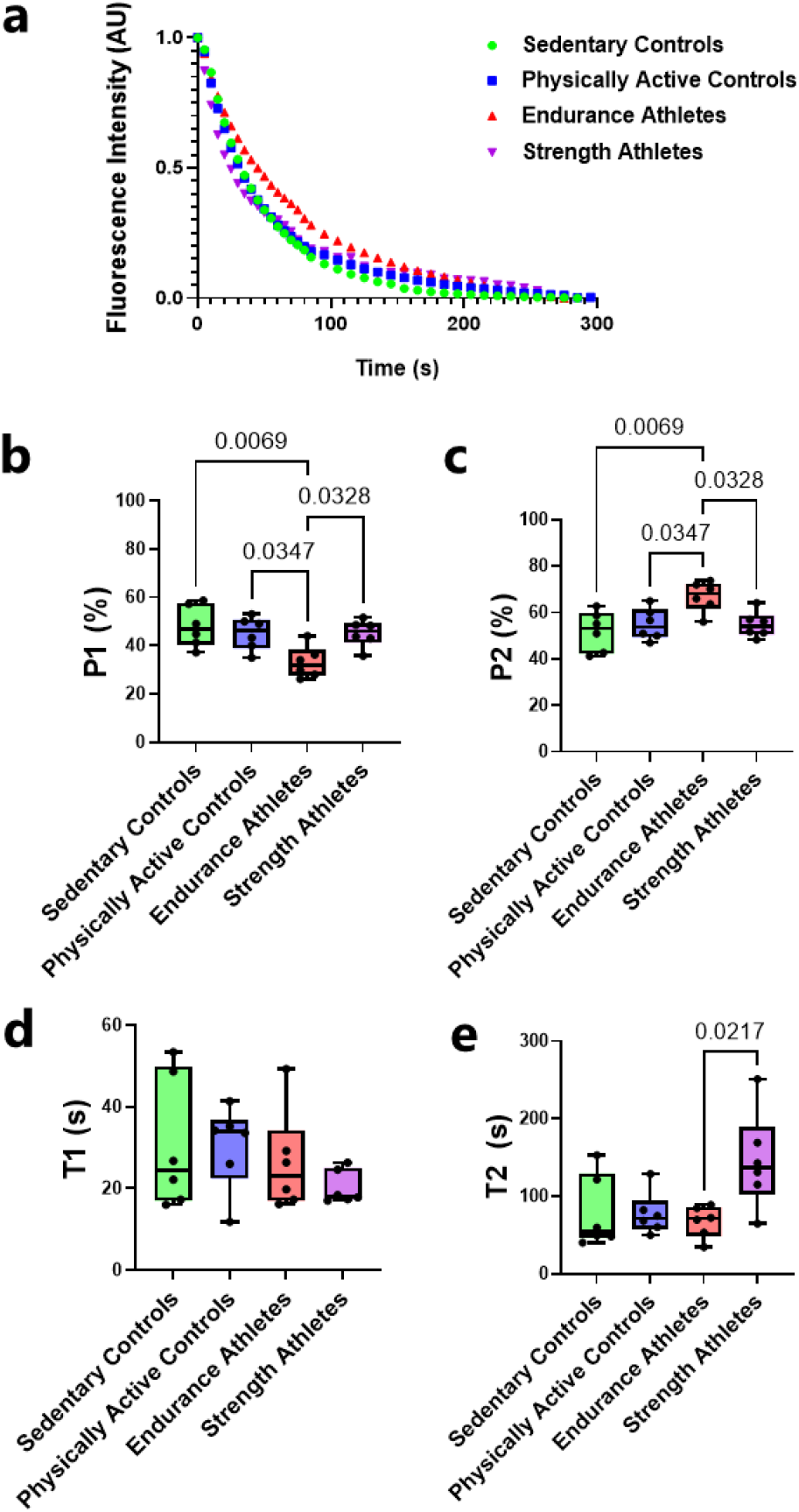
Myosin head conformation and ATP turnover time is changed in athlete skeletal myofibres. (**a**) A representative Mant-ATP chase experiment decay graph showing exponential decays of single muscle fibres from sedentary controls, physically active controls, endurance athletes and strength athletes. The Percentage of myosin heads in skeletal myofibres in the (**b**) P1 denoting the disordered-relaxed state (DRX) and (**c**) P2 denoting the super-relaxed state (SRX). This was estimated from the equation shown in the methods section. (**d**) T1 value in seconds denoting the ATP turnover lifetime of the DRX. (**e**) T2 value in seconds denoting the ATP turnover lifetime in the SRX. Each data point represents the mean value from all fibres from of each subject. Data is presented as a box and whiskers plot with min to max shown on error bars. Five individual muscle fibres were analyzed from a total of six subjects per group.

### Minimal effects of Mavacamten on SRX myosins in endurance athletes

As we only observed significant differences in the amount of SRX myosins in the EA group, the following experiments focused exclusively on this particular group of individuals. We wanted to further test whether Mavacamten, a new FDA approved SRX-inducing drug for the treatment of HCM, would be able to be potent in individuals where the amount of SRX myosins is already high (Figure 2c). Mavacamten is known to specifically target the β/slow-cardiac myosin (Anderson et al., 2018, Heitner et al., 2019). Hence, we only investigated the effects of the drug on skeletal myofibres expressing this β/slow-cardiac myosin. We did not find any significant change in the proportions of DRX or SRX myosins following 0.3 μM Mavacamten treatment (Figure 3a and b). Furthermore, the ATP turnover time of DRX myosins was unchanged following mavacamten treatment (Figure 3c). We did however observe significantly prolonged SRX myosin ATP turnover lifetime following treatment with 0.3 μM Mavacamten (Figure 3d).

**Figure 3.**
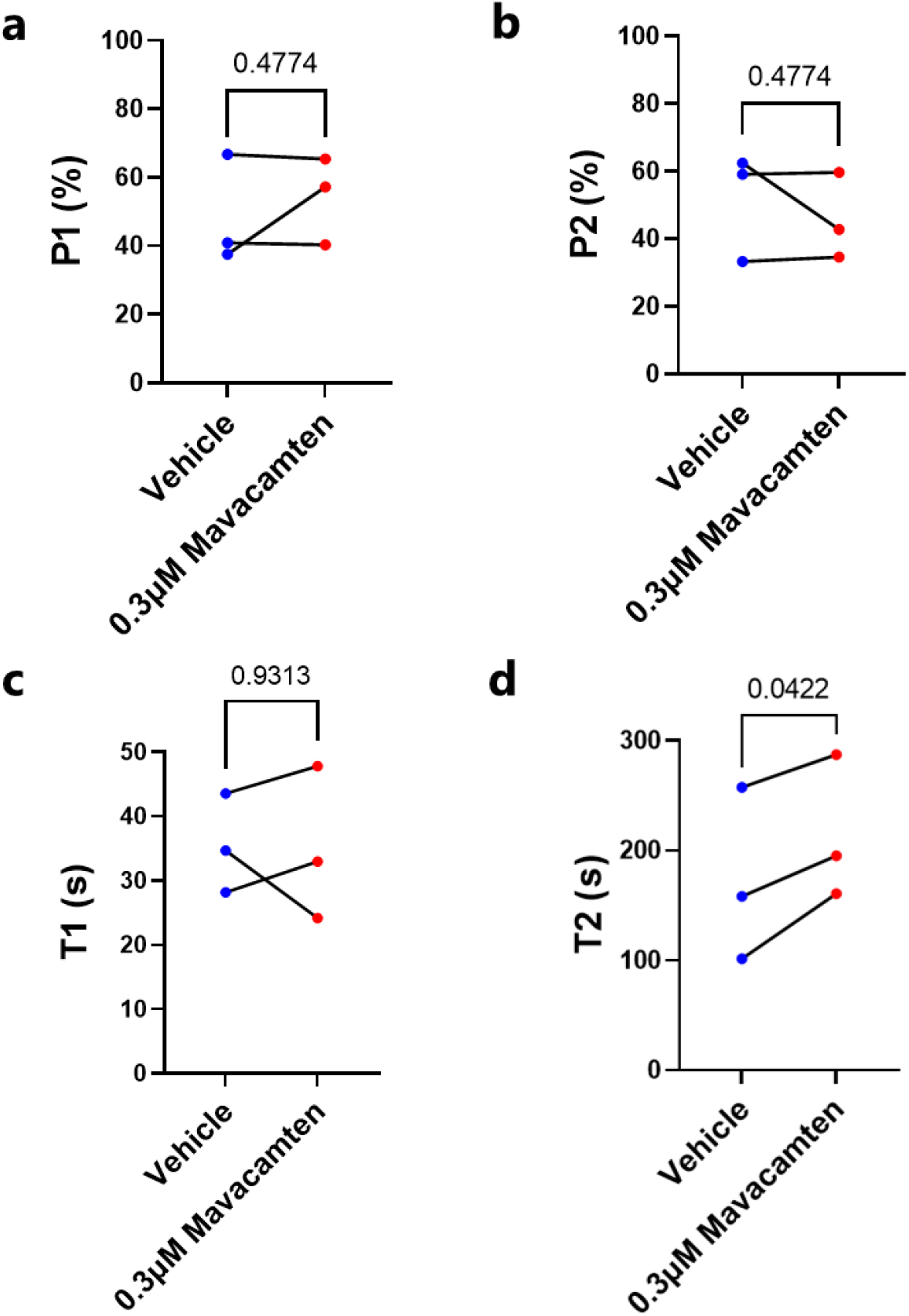
Mavacamten treatment does not influence myosin head conformation percentages in type I but increases SRX ATP turnover lifetime in skeletal myofibres of endurance athletes. (**a**) P1 denoting disordered-relaxed state (DRX) and (**b**) P2 denoting the super-relaxed state (SRX) before and after treatment with 0.3 μM Mavacamten. (**c**) T1 in seconds denoting the ATP turnover time of the disordered-relaxed state (DRX) before and after 0.3 μM Mavacamten treatment. (**d**) T2 in seconds denoting the ATP turnover time of the super-relaxed state (SRX) before and after treatment with 0.3 μM Mavacamten. The mean value of 6 fibres from each subject are shown paired as before and after Mavacamten treatment. This was estimated from the equation shown in the methods section. In panel (d), single muscle fibre values of greater than 500s were excluded as outliers. n=3. p-values are shown on each panel.

## Discussion

In this study, we, for the first time, examined to which extent myosin head conformations in relaxed human skeletal myofibres differ between highly trained athlete populations compared to recreationally active and sedentary controls. Our data indicate that endurance-trained athletes have higher proportions of SRX myosins, and that Mavacamten is not responsive in this specific population. Overall, these suggest that myosin head conformations can be modulated by changes in the functional or environmental demand and that drugs targeting myosin heads may not be efficient in extremely physically active populations.

Physical activity is well known to have vast whole-body metabolic benefits including, but not limited to, the improvement of mitochondrial functional, the removal of reactive oxygen species, re-wiring of insulin signaling in skeletal muscle and the prevention of obesity (Holten et al., 2004, Needham et al., 2022, Richter and Hargreaves, 2013, Memme et al., 2021). However, how different types of physical exercise background may influence the basal metabolic rate of skeletal muscle has, until now, not been studied. Different training regimes have been previously shown effect the biology of skeletal muscle via such switching of the myosin heavy chain (MyHC) expression, and thus fibre-type within skeletal muscle (Aagaard and Andersen, 1998). In particular, an increase in MyHC IIA expression, denoting fast muscle fibres, is associated with strength training and an increase in the maximal force and power which is able to be produced by a muscle (Harridge et al., 1996). Whereas, a preferential expression of MyHC I and thus a higher proportion of slow, oxidative fibres, is associated with endurance training (Andersen and Aagaard, 2010). In the present study, myosin conformational states and their fine-tuning are highlighted as potential regulators of muscle metabolism. Importantly, in the present study, we observed that in EA, the percentage of myosin heads in the SRX state was significantly greater when compared to SC, PAC and SA subjects, suggesting chronic endurance training may decrease the overall resting metabolic rate of the skeletal muscle. This may allow for a maximum supply of ATP for utilization during exercise or optimal ATP recovery post-exercise or between successive interval exercise bouts. Specific Further studies into the fibre-type specific resting myosin regulation would further benefit this field going forward. As it is well documented that increased levels of physical fitness are associated with a lower resting heart-rate and increased stroke-volume, it would be of interest to observe if highly-trained individuals have altered cardiac SRX myosins as well, or if the myosin head conformation is linked to heart rate plasticity or cardiac output in any manner (Cornelissen et al., 2010).

Mechanisms underlying the exercise–induced increase in SRX myosins remain unknown, however, previous studies have demonstrated the importance of post-translational modifications in the regulation of the myosin function (Alamo et al., 2016, Brito et al., 2011, McNamara et al., 2019a, McNamara et al., 2019b). It has been established that exercise can induce vast protein phosphorylation, including that of sarcomeric proteins, and that different training methods can induce distinct training-specific phosphorylation profiles (Needham et al., 2022, Bódi et al., 2021). Further research into the PTMs that specifically occur in endurance-trained individuals could provide insight to why we see this result in EAs and into the mechanisms of control of myosin head conformation.

The approval of Mavacamten for the treatment of HCM has highlighted the potential of targeting resting myosin heads for the treatment of both cardiac and skeletal muscle diseases. This strategy is particularly promising due to the elusiveness of treating myopathies to date. Although already FDA-approved the exact mechanism of action of Mavacamten is not fully understood. Studies have shown, that Mavacamten can stabilize the SRX state within cardiac myocytes, thus decreasing the percentage of pathological DRX myosins in HCM (Olivotto et al., 2020, Anderson et al., 2018). Here, we did not find any change in the percentages of SRX myosins following Mavacamten treatment at 0.3 μM. This result suggests that Mavacamten may not be as potent in skeletal muscle fibres from chronically trained individuals as in cardiac myocytes from patients. One plausible explanation may lie in the already high level of SRX myosins in the endurance athlete population. Other possible causes may relate to differences between skeletal and cardiac tissue. Indeed, although the core mechanism of the resting myosin conformations are similar between skeletal and cardiac muscle, they do differ functionally in their behavior upon contraction. Upon the contraction of skeletal muscle, all myosin heads, regardless of their conformation at rest are recruited to bind to the actin thin-filament and facilitate contraction (Hooijman et al., 2011). In cardiac muscle however, myosin heads in the SRX state remain in this state during contraction and do therefore do not form interactions with actin (Schmid and Toepfer, 2021). These differences point to a core difference in the stability of the SRX conformation between skeletal and cardiac muscle cells and thus a possible explanation to the lack of resting myosin conformation change following Mavacamten treatment in our study. Studies have also previously shown that the affinity of Mavacamten to skeletal muscle in rabbit skinned skeletal muscle fibres is far lower than cardiac fibres, however this has yet to be confirmed in humans (Green et al., 2016). Further mechanistic studies have suggested that Mavacamten primarily reduces the steady-state ATPase activity by inhibiting the rate of phosphate release of β/slow myosin (Kawas et al., 2017). Our results are somewhat consistent with this finding as we demonstrate that Mavacamten slows the ATP turnover lifetime of SRX myosins, perhaps indicating a further stabilization of the SRX myosin state following Mavacamten treatment in skeletal muscle. This means that Mavacamten can slow the rate of ATP consumption in skeletal muscle fibres from endurance athletes, without changing the actual percentage of myosin heads in a certain conformation.

In summary, the present study indicates that resting myosin conformation is influenced by training status and that high-level endurance trained subjects had a greater percentage of SRX myosin heads in skeletal muscle and thus a potentially reduced resting metabolic rate in their skeletal muscle. Additionally, we show that, Mavacamten, known to also increase the amount of SRX myosins in cardiac muscle, is not efficient in the skeletal muscle of highly endurance-trained individuals. All these findings are important as they suggest a dynamic plasticity of resting myosin conformations linked to exercise that may not be entirely compatible with myosin-driven drug treatments.

## Author Contributions

**CTAL**: Conceived Study, performed experiments, analyzed data, interpreted results, wrote manuscript, managed project. **LT**: Performed experiments, analyzed data, interpreted results, wrote manuscript. **JL and TNB**: performed experiments, critically reviewed manuscript. **JN and PA**: Conceived study, provided subject samples, critically reviewed manuscript. **JLA, CS, JWH, FD, SL, RES, RH, SiL, AI, TR, MTH and CS**: Provided subject samples, critically reviewed manuscript. **JO**: Conceived study, interpreted results, wrote manuscript, managed project, acquired funding. All authors approved the final version of the manuscript.

